# Mechanical Profiling of Biopolymer Condensates through Acoustic Trapping

**DOI:** 10.1101/2024.09.16.613217

**Authors:** Kichitaro Nakajima, Tomas Sneideris, Lydia L. Good, Nadia A. Erkamp, Hirotsugu Ogi, Tuomas P. J. Knowles

## Abstract

Characterizing the mechanical properties of single colloids is a central problem in soft matter physics. It also plays a key role in cell biology through biopolymer condensates, which function as membraneless compartments. Such systems can also malfunction, leading to the onset of a number of diseases, including many neurodegenerative diseases; the functional and pathological condensates are commonly differentiated by their mechanical signature. Probing the mechanical properties of biopolymer condensates at the single particle level has, however, remained challenging. In this study, we demonstrate that acoustic trapping can be used to profile the mechanical properties of single condensates in a contactless manner. We find that acoustic fields exert the acoustic radiation force on condensates, leading to their migration to a trapping point where acoustic potential energy is minimized. Furthermore, our results show that the Brownian motion fluctuation of condensates in an acoustic potential well is an accurate probe for their bulk modulus. We demonstrate that this framework can detect the change in the bulk modulus of polyadenylic acid condensates in response to changes in environmental conditions. Our results show that acoustic trapping opens up a novel path to profile the mechanical properties of soft colloids at the single particle level in a non-invasive manner with applications in biology, materials science, and beyond.

## 1 Introduction

Characterizing the mechanical properties of soft colloids is a key issue in soft matter physics, and its significance has recently been highlighted in cell biology by the emergence of biopolymer condensates composed of proteins and/or nucleic acids ^1^. These condensates play crucial roles in regulating a range of physiological functions within cells ^1–3^. They exhibit liquid-like properties, allowing them to dynamically assemble and disassemble in response to cellular signals ^4,5^. On the other hand, certain proteins have been shown to form liquid-like condensates that can undergo an aging process, transitioning into a more solid-like state, which has been implicated in the pathology of neurodegenerative diseases ^6–8^. The aging of condensates is typically associated with a loss of dynamic behavior of biopolymers within the condensate and an increase in elastic moduli ^9,10^. Therefore, a tool that can accurately measure the mechanical properties of biopolymer condensates at the single-particle level is crucial for understanding the molecular mechanisms behind both functional and pathological condensates.

The material properties of biopolymer condensates have been investigated using various analytical techniques ^11,12^, including micropipette aspiration ^13^, microrheology ^14,15^, and coalescence assays ^16^. These methods have successfully characterized their viscous fluid properties, such as viscosity and surface tension. However, accurately assessing their mechanical properties as elastic bodies, including elastic modulus and bulk modulus, remains a significant challenge. This difficulty arises primarily from the challenge of applying external forces to a single mesoscale condensate on timescales adequately shorter than its relaxation time of deformation (typically 10 ms − 1 s ^8,17^), where viscoelastic materials predominantly exhibit elastic behavior.

Here, we focus on acoustic trapping as an analytical tool for characterizing the mechanical properties of biopolymer condensates. Acoustic trapping ^18,19^, analogous to optical trapping ^20,21^, utilizes the acoustic radiation force ^22^, which acts on microparticles within a standing pressure field ^22^. The acoustic radiation force arises from the interference between an elastic body in the acoustic field and ultrafast pressure oscillations of the acoustic wave ^19,22^, whose timescale is significantly shorter than the relaxation time of the condensates. Consequently, acoustic trapping potentially provides a unique capability to measure the elastic nature of mesoscale soft colloids at the single particle level. Moreover, acoustic trapping is inherently biocompatible, as it does not involve focused laser irradiation or the use of beads. This bio-compatibility has enabled the manipulation of cells ^23,24^, extra-cellular vesicles ^25^, and embryos ^26^ without compromising their viability ^27^. However, acoustic trapping has yet to be applied in research on the mechanical properties of biopolymer condensates.

In this study, we fabricate acoustic devices composed of a microfluidic channel and piezoelectric substrate to generate surface acoustic waves (SAWs) with frequencies of ∼50 MHz, thereby creating an acoustic potential field in the sample solution containing condensates (**Fig. 1a,b**). The acoustic potential field exerts an acoustic radiation force on the condensates (**Fig. 1b**), allowing us to assess their material properties. We show that polyadenylic acid (poly-rA) condensates can be trapped with a trapping stiffness of ∼100 fN/µm and aligned into a lattice pattern corresponding to the acoustic potential field in a contactless manner. By trapping two condensates in a single acoustic potential well, we can induce their coalescence events to analyze the inverse capillary velocity ^4^. We present a novel framework focusing on the positional fluctuations of the Brownian motion of condensates as an accurate probe of their bulk modulus. We use this approach to investigate the change in the bulk modulus of poly-rA condensates in response to environmental conditions.

**Fig. 1.**
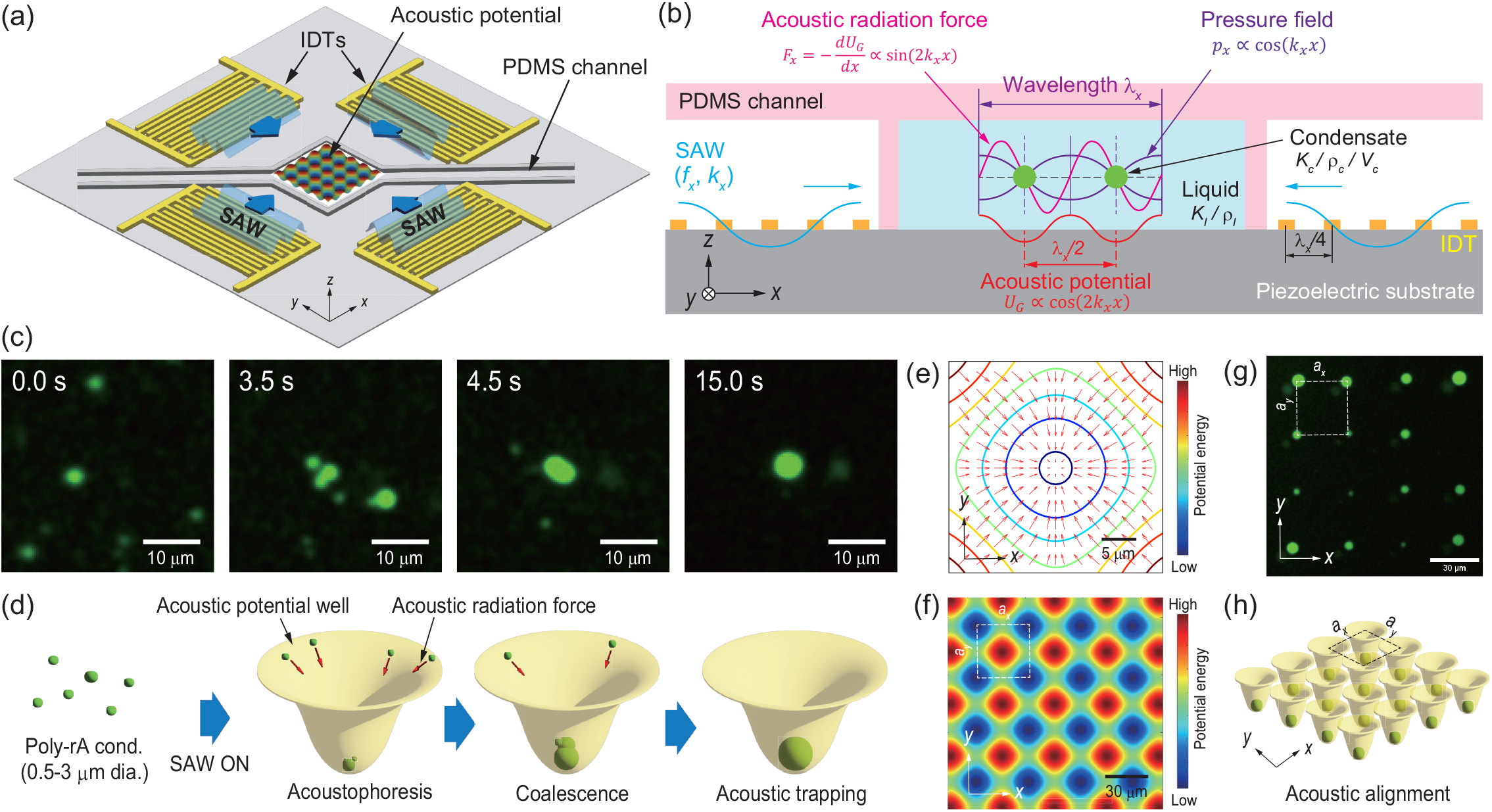
Acoustic trapping of biopolymer condensates. (a) Schematic illustration of the acoustic device developed in this study. By applying an alternative voltage to interdigital transducers (IDTs), IDTs emit surface acoustic waves (SAWs). The SAWs create a two-dimensional (2D) acoustic potential within the PDMS microfluidic chamber. (b) Working principle of the acoustic device for trapping condensates. The SAWs produce a standing acoustic-pressure field *p*_*x*_ in the sample solution, resulting in the acoustic potential field *U*_*G*_ and an acoustic radiation force *F*_*x*_ that acts on condensates. (c) Fluorescent images (Movie S2) and (d) schematic illustration of poly-rA condensates trapped by acoustic radiation force. Scale bars: 10 µm. (e) Acoustic potential (colored lines) and acoustic radiation force (red arrows) in the 2D acoustic field with *f*_*x,y*_ = 49.6*/*49.7 MHz. Scale bar: 5 µm. (f) Calculated acoustic potential field formed by the 2D acoustic field with *f*_*x,y*_ = 49.6*/*49.7 MHz. Scale bar: 30 µm. (g) Experimental fluorescence image and (h) schematic illustration of poly-rA condensates aligned by the 2D acoustic field with *f*_*x,y*_ = 49.6*/*49.7 MHz. Scale bar: 30 µm.

## 2 Results

### 2.1 Acoustic trapping of biopolymer condensates

We constructed an acoustic device composed of a 128^°^ Y-cut lithium niobate (LN) substrate and a polydimethylsiloxane (PDMS) microfluidic channel. The PDMS channel possesses a square chamber with a volume of 500 × 500 × 53 µm^3^. The LN substrate features two pairs of interdigital transducers (IDTs) that are orthogonal to each other. By positioning the PDMS chamber at the intersection of these two pairs of IDTs and applying an alternating voltage to the IDTs (**Fig. 1a** and **Fig. S1**), a 2D standing pressure field can be generated in the sample solution.

When a particle with a bulk modulus and density different from those of the surrounding liquid medium is placed in a standing pressure field, an acoustic radiation force is exerted on the particle (**Fig. 1b**). Theoretically, the acoustic radiation force *F* acting on a spherical particle in an inviscid fluid can be expressed as the gradient of the Gor’kov acoustic potential *UG* ^28,29^ given by

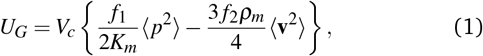

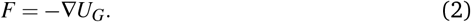

Here, 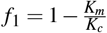 and 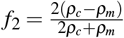, where *K*_*m*_ and *K*_*c*_ are the bulk moduli of the liquid medium and the particle (or condensate), respectively, and *ρ*_*m*_ and *ρ*_*c*_ are their mass densities. *V*_*c*_ denotes the volume of the particle. ⟨*p*^2^⟩ and ⟨**v**^2^⟩ are the time-averaged pressure and velocity field squared, respectively.

In the one-dimensional (1D) case (**Fig. 1b**), the SAWs emitted from a pair of the IDTs along the *x*-axis create a standing pressure field *p*_*x*_ = *A*_*x*_cos(*k*_*x*_*x*)cos(2*π f*_*x*_*t*) in the liquid medium. Here, *A*_*x*_, *k*_*x*_, and *f*_*x*_ are the amplitude of the pressure field, the wavenumber, and the frequency of the SAW along the *x*-axis, respectively. Under this pressure field, the velocity field is derived as 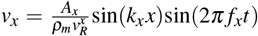 sin(*k*_*x*_*x*)sin(2*π f*_*x*_*t*) using the relationship 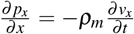, where 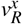 denotes the phase velocity of the SAW propagating along the *x*-axis. By substituting *p*_*x*_ and *v*_*x*_ into **Eqs. 1** and **2**, we obtain the acoustic potential and acoustic radiation force along the *x*-axis in the 1D case as

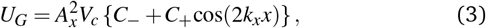

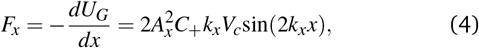

where 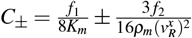. For soft colloids dispersed in an aqueous solution, *C*_+_ is typically positive, as the density and bulk modulus of the particle are higher than those of the liquid medium (*ρ*_*c*_ *> ρ*_*m*_ and *K*_*c*_ *> K*_*m*_) ^30^. This results in the migration of the particles, a phenomenon known as acoustophoresis ^31^, to the nodes. Consequently, microparticles are aligned along the *x*-axis with a spacing equal to half the wavelength of the SAW (**Fig. 1b**).

The working principle of acoustic trapping in the 2D case is essentially the same as in the 1D case. By introducing a second standing pressure field along the *y*-axis, which is perpendicular to the *x*-axis, given by *p*_*y*_ = *A*_*y*_cos(*k*_*y*_*y*)cos(2*π f*_*y*_*t*), we can calculate the 2D acoustic potential field. Here, *A*_*y*_, *k*_*y*_, and *f*_*y*_ represent the amplitude of the pressure field, the wavenumber, and the frequency of the SAW along the *y*-axis, respectively. By using ⟨*p*^2^⟩ = ⟨(*p*_*x*_ + *p*_*y*_)^2^⟩ and 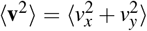, we can derive the time-averaged pressure and velocity field squared as

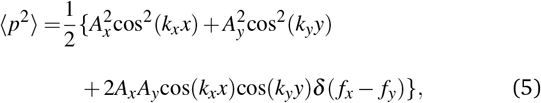

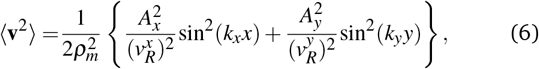

where *δ* (·) is the Kronecker’s delta function, and 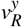denotes the phase velocity of the SAW propagating along the *y*-axis. The 2D acoustic potential and acoustic radiation force can be calculated by substituting **Eqs. 5** and **6** into **Eqs. 1** and **2**. We experimentally confirmed that the 2D acoustic field predicted theoretically is formed in our experimental setup using polystyrene particles with an average diameter of 1.0 *µ*m (**Fig. S2** and **Movie S1**).

In this study, we employed poly-rA condensates as a biopolymer condensate. Upon the addition of sufficient NaCl, poly-rA undergoes phase separation into dense condensates. Under the experimental conditions, condensates typically range from 0.5 to 3 µm in diameter (**Fig. 1c**). By generating a 2D standing acoustic field, the dispersed poly-rA condensates migrate towards a trapping point via acoustic radiation force (**Fig. 1c,d** and **Movie S2**). This migration results in the coalescence of individual condensates into a larger, single condensate, which remains trapped. This observation shows that the acoustic field exerts the acoustic radiation force on the poly-rA condensates, driving them towards the trapping point where the acoustic potential energy is minimized (**Fig. 1e**).

**Figure 1f** depicts the theoretical acoustic potential field generated by the 2D acoustic pressure field with frequencies of *f*_*x,y*_ = 49.6*/*49.7 MHz. The acoustic potential exhibits a lattice structure with a lattice constant of *a*_*x,y*_ ∼ 36.1 µm. In the experiment, a similar lattice structure of poly-rA condensates was observed, with lattice constants of *a*_*x,y*_ = 34.2 ± 1.3*/*34.2 ± 1.6 µm (**Fig. 1g,h**). The experimentally determined lattice constant was in close agreement with the theoretical value, providing robust evidence that the acoustic radiation force can simultaneously trap multiple biomolecular condensates in a contactless manner.

### 2.2 Acoustically-induced coalescence of condensates

The inverse capillary velocity, which quantifies the ratio between the surface tension and viscosity of the condensates ^16^, can be analyzed by examining the kinetics of the coalescence events for condensates with various diameters. This parameter is essential for evaluating the liquid-like properties of biomolecular condensates ^32^ and is commonly employed to monitor the transition from liquid-like to solid-like states as the condensates age ^33^.

In the experiment, two poly-rA condensates initially separated by approximately 20 µm were brought together at a single trapping point, where they subsequently merged into a single condensate within *t* = 60 s (**Fig. 2a,b** and **Movie S3**). The time-course plot of the aspect ratio of the condensate during the coalescence event demonstrates an exponential relaxation behavior (**Fig. 2c**). The experimental data are well-fitted with the function *f* (*t*) = *A*·exp(− *t/τ*) + 1, where *A* is a fitting parameter, and *τ* denotes the relaxation time ^11^. These results indicate that acoustic trapping does not bias the intrinsic kinetics of the coalescence.

**Fig. 2.**
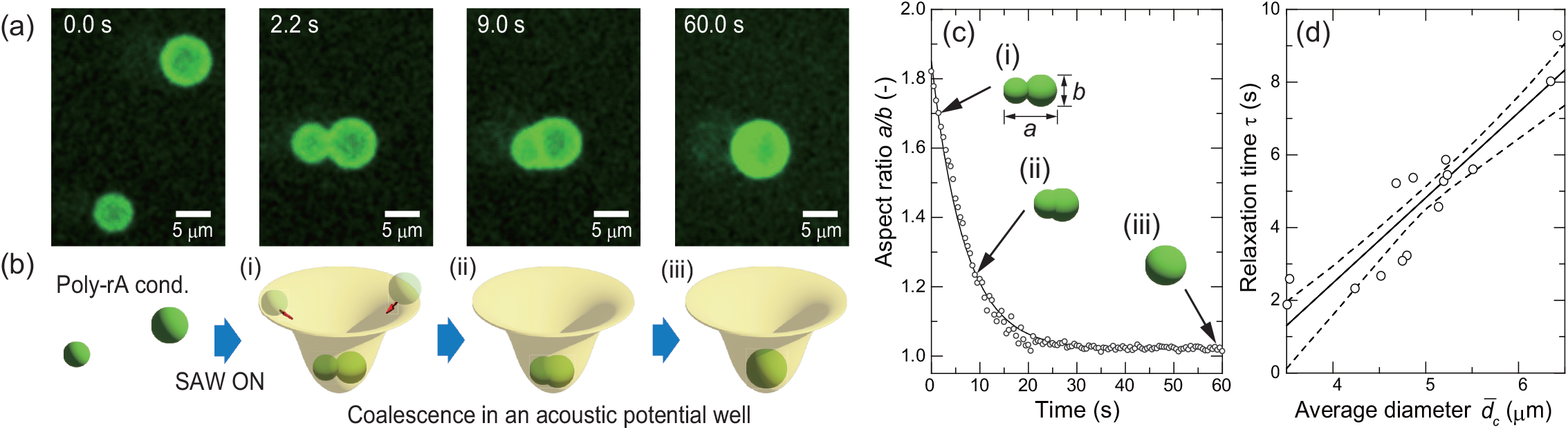
Analysis of the inverse capillary velocity by acoustic trapping. (a) Fluorescent images (Movie S3) and (b) schematic illustration of the acoustically-induced coalescence event between two poly-rA condensates. Scale bars: 5 µm. (c) Time-course plots of the aspect ratio of poly-rA condensates (circles) during the coalescence event, with a fitted curve to determine the relaxation time (solid curve). (d) Relaxation times of coalescence for poly-rA condensates of different average diameters (*n* = 15). The solid and dotted lines represent an average of the linear fit and its 95% confidence interval, respectively. The slope indicates the inverse capillary velocity of poly-rA condensates, *η*_*c*_*/γ*_*c*_ = 2.34 ± 0.27 s/µm. The poly-rA condensates are formed with [Poly-rA] = 1.0 mg/mL, [NaCl] = 1.0 M, and [HEPES] = 50 mM.

The relaxation time of coalescence, *τ*, can be expressed as 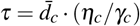, where 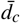 is the average diameter of the two condensates, and *η*_*c*_ and *γ*_*c*_ represent the viscosity and surface tension of the condensates, respectively ^4^. This relationship allows for determining the inverse capillary velocity, *η*_*c*_*/γ*_*c*_. **Figure 2d** presents the correlation between the relaxation time and the average diameter for fifteen independent coalescence events. The linear fit shows a good correlation with the experimental data, yielding a Pearson correlation coefficient of |*R*| = 0.92. We calculated the inverse capillary velocity from this fit to be 2.34 ± 0.27 s/µm for poly-rA condensates. These results demonstrate that acoustic trapping can effectively induce the coalescence of condensates by bringing two condensates together at a trapping point, facilitating the analysis of the inverse capillary velocity.

### 2.3 Brownian motion of acoustically-trapped condensates

The condensates trapped within the acoustic field exhibit the confined Brownian motion in the acoustic potential well. By experimentally tracking the trajectory of this Brownian motion (**Fig. 3a**), we quantified its positional fluctuations, *σ*, by measuring the standard deviation of the spatial distribution of the condensate centers. An increase in the applied voltage to the IDT resulted in more pronounced confinement of the Brownian motion of the poly-rA condensates and thus smaller positional fluctuations (**Fig. 3b**). The acoustic trapping stiffness on poly-rA condensates can be determined using the equipartition theorem of energy ^34^, given by *κ* = 2*k*_*B*_*T/σ* ^2^, where *κ, k*_*B*_, and *T* denote the trapping stiffness, the Boltzmann constant, and the absolute temperature, respectively. For poly-rA condensates with a diameter of 3.42 ± 0.04 µm, the trapping stiffness is on the order of 10 to 100 fN/µm (**Fig. 3b**).

**Fig. 3.**
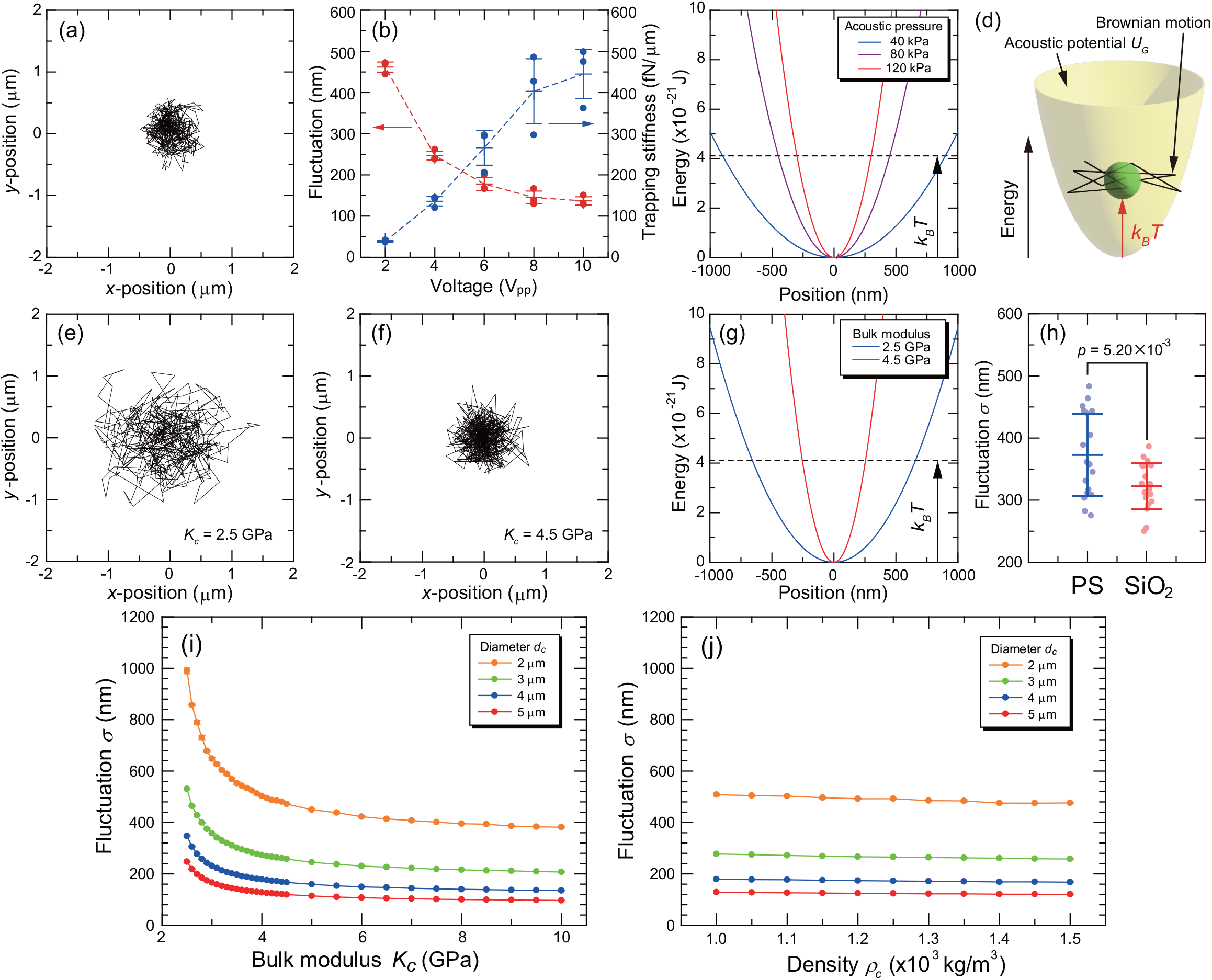
Brownian motion of a single condensate in an acoustic potential well. (a) Experimental trajectory of the Brownian motion of the poly-rA condensate with a diameter of 3.42 ± 0.04 µm trapped by acoustic frequencies of *f*_*x,y*_ = 35.2*/*35.3 MHz and the applied voltage of 4 V_pp_. (b) Fluctuation of the Brownian motion of the acoustically-trapped poly-rA condensates with a diameter of 3.42 ± 0.04 µm (red plots) and trapping stiffness (blue plots) calculated from the fluctuation with various voltages applied on the IDTs. The acoustic frequencies are *f*_*x,y*_ = 35.2*/*35.3 MHz. The poly-rA condensates are formed with [Poly-rA] = 1.0 mg/mL, [NaCl] = 1.0 M, and [HEPES] = 50 mM. Error bars denote the standard deviation among three independent measurements. (c) Computed acoustic potentials under various acoustic pressures. Simulation parameters: *A*_*x,y*_ = 40*/*80*/*120 kPa, *f*_*x,y*_ = 35.2*/*35.3 MHz, *d*_*c*_ = 3.0 µm, *K*_*c*_ = 4.55 GPa, *ρ*_*c*_ = 1, 120 kg/m^3^, *ρ*_*m*_ = 997 kg/m^3^, *K*_*m*_ = 2.22 GPa, and *η*_*m*_ = 8.94 × 10^−4^ Pa·s. (d) Schematic illustration of the Brownian motion of a condensate in an acoustic potential well, determined by the interplay between the thermal energy *k*_*B*_*T* and acoustic potential *U*_*G*_. (e,f) Trajectories of the Brownian motion of acoustically trapped condensates with bulk moduli of (e) *K*_*c*_ = 2.5 GPa and (f) *K*_*c*_ = 4.5 GPa, from the Langevin dynamics simulation. (g) Computed acoustic potentials for condensates with various bulk moduli. Simulation parameters: *A*_*x,y*_ = 100 kPa, *f*_*x,y*_ = 35.2*/*35.3 MHz, and *d*_*c*_, *ρ*_*c*_, *ρ*_*m*_, *K*_*m*_, and *η*_*m*_ are the same as in panel c. (h) Experimental fluctuations of Brownian motion in acoustic potential wells for polystyrene (PS) and silica (SiO_2_) microspheres with diameters of 2.26 ± 0.16 µm and 2.21 ± 0.10 µm, respectively. Experimental conditions: *A*_*x,y*_ = 100 kPa, *f*_*x,y*_ = 35.2*/*35.3 MHz. Error bars represent the standard deviation among independent measurements (*n*=18). (i,j) Dependencies of fluctuation on (i) bulk modulus and (j) density of condensates. Simulation parameters; *ρ*_*c*_ = 1, 120 kg/m^3^ (for panel i), *K*_*c*_ = 4.0 GPa (for panel j), and *A*_*x,y*_, *f*_*x,y*_, *ρ*_*m*_, *K*_*m*_, and *η*_*m*_ are the same as in panel g.

The trapping stiffness corresponds to the curvature of the acoustic potential near its minima. The higher the applied voltage to the IDTs, the greater the amplitude of the acoustic field ^22^, resulting in the steeper curvature of the potential in proximity to the minima (**Fig. 3c**). The magnitude of this Brownian motion fluctuation is governed by the interplay between thermal energy, *k*_*B*_*T*, and the potential energy created by the standing acoustic waves, *U*_*G*_ (**Fig. 3d**). Thus, the more pronounced confinement of the Brownian motion of the condensates in the acoustic potential well can be understood by the steeper curvature of the acoustic potential at the higher acoustic pressure.

In theory, the Brownian motion of a micrometer-sized condensate in an acoustic potential well can be described by the overdamped Langevin equation with an acoustic potential term ^28,29,35^,

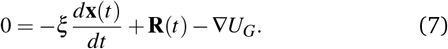

Here, **x**(*t*) represents the position of the center of the condensate, and **R**(*t*) denotes the stochastic force corresponding to thermal fluctuation. The friction constant *ξ* is given by *ξ* = 3*πη*_*m*_*d*_*c*_, where *d*_*c*_ and *η*_*m*_ are the diameter of the condensate and the viscosity of the liquid medium, respectively. The stochastic force **R**(*t*) satisfies the relationships ⟨**R**(*t*)⟩ = 0 and ⟨**R**(*t*) · **R**(*t*′)⟩ = 4*ξk*_*B*_*Tδ* (*t* − *t*′).

It is noteworthy that the acoustic potential includes a term related to the bulk modulus of the trapped condensate (**Eq. 1**), suggesting that the bulk modulus influences the Brownian motion of the condensate. We conducted Langevin dynamics simulations of Brownian motion trajectories for acoustically trapped condensates with a diameter of *d*_*c*_ = 3 *µ*m, varying their bulk moduli while keeping other parameters constant (**Figure 3e,f**). The simulated trajectories for condensates with bulk moduli of *K*_*c*_ = 2.5 and 4.5 GPa, respectively, under identical acoustic fields. The condensate with the higher bulk modulus (**Fig. 3f**) exhibits less fluctuation compared to the one with the lower bulk modulus (**Fig. 3e**). These results reflect the steeper curvature of the acoustic potential on the stiffer condensate than the softer one (**Fig. 3g**). The acoustic radiation force results from the interference between the incoming wave and the wave scattered by the particle in the acoustic field ^22^. Stiffer condensates scatter the incoming waves more significantly than softer ones due to the greater mismatch in bulk modulus between the condensate and the liquid medium. This increased amplitude of the scattered wave enhances the acoustic radiation force acting on the stiffer condensates, leading to more confined Brownian motion in the acoustic potential well.

To experimentally validate the computational results, we conducted model experiments using polystyrene (PS) and silica (SiO_2_) particles. The fluctuations of the Brownian motion of these particles were observed in the acoustic potential well created by an acoustic field with amplitudes and frequencies of *A*_*x,y*_ = 100 kPa and *f*_*x,y*_ = 35.2*/*35.3 MHz (**Fig. 3h**). Here, the acoustic pressures *A*_*x,y*_ were calibrated through a separate experiment (**Fig. S3**). Although there was no statistically significant difference in the diameters of the particles (2.26 ± 0.16 *µ*m for PS and 2.21 ± 0.10 *µ*m for SiO_2_), a significant difference in the fluctuations was observed. The density and bulk modulus of SiO_2_ (*ρ*_*c*_ = 2.20 kg/m^3^, *K*_*c*_ = 36.3 GPa ^36^) are higher than those of PS (*ρ*_*c*_ = 1.06 kg/m^3^, *K*_*c*_ = 4.55 GPa ^30^). The observed smaller fluctuation in SiO_2_ particles compared to PS particles suggests that the fluctuation of the Brownian motion reflects their mechanical properties, which are detectable in our experimental setup.

**Figure 3i** illustrates the relationship between fluctuation and the bulk modulus of condensates with various diameters. We focus on this dependency to analyze the bulk moduli of poly-rA condensates based on their Brownian motion in an acoustic potential well. Notably, fluctuations are highly sensitive to changes in the bulk modulus within the range of *K*_*c*_ = 2.5−4.0 GPa. The acoustic radiation force acting on the condensate is determined by the relative difference between the bulk modulus of the condensate and that of the liquid medium. Consequently, fluctuations show significant sensitivity to the bulk modulus when it is close to that of the liquid medium (*K*_*m*_ = 2.2 GPa for water). Conversely, when the bulk modulus of the condensate deviates substantially from that of the liquid medium, fluctuations become less sensitive to further changes.

Two additional parameters that possibly influence the fluctuation of Brownian motion of acoustically trapped condensates are the density of the condensate, *ρ*_*c*_, and the viscosity of the liquid medium, *η*_*m*_. Langevin dynamics simulations show that fluctuations remain relatively unchanged as the density of the condensates varies (**Fig. 3j**). Although an increase in density generally leads to a decrease in fluctuation, this effect is much smaller than that of the bulk modulus because the first term on the right-hand side of **Eq. 1** is dominant over the second term. Therefore, for the subsequent analysis, the second term on the right-hand side of **Eq. 1** is treated as a constant, using *ρ*_*c*_ = 1,120 kg/m^3^, the density reported for germinal vesicles ^37^. Since the Gor’kov acoustic potential (**Eq. 1**) is derived under the assumption that the liquid medium is inviscid, the results are unaffected by viscosity in our simulation model. It has been reported that when the thickness of the viscous boundary layer 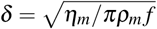 is much smaller than the diameter of the trapped particles, the effects of viscosity are negligible ^38^. In our experimental conditions, the *δ* value is less than 100 nm, so the effects of viscosity on the acoustic radiation force and resulting Brownian motion are insignificant.

### 2.4 Analysis of bulk moduli of condensates

Condensates with different bulk moduli show different dependencies of fluctuation on their diameters (**Fig. 4a**). We focused on this relationship to experimentally determine the bulk moduli of poly-rA condensates. We measured the fluctuations in Brownian motion for condensates with various diameters and compared these experimental results with those obtained from simulations. Then, the mean square errors (MSEs) between the experimental and simulated results were calculated across a range of bulk moduli. The bulk modulus that minimized MSE was determined to be the bulk modulus of the condensates (**Fig. 4b**).

**Fig. 4.**
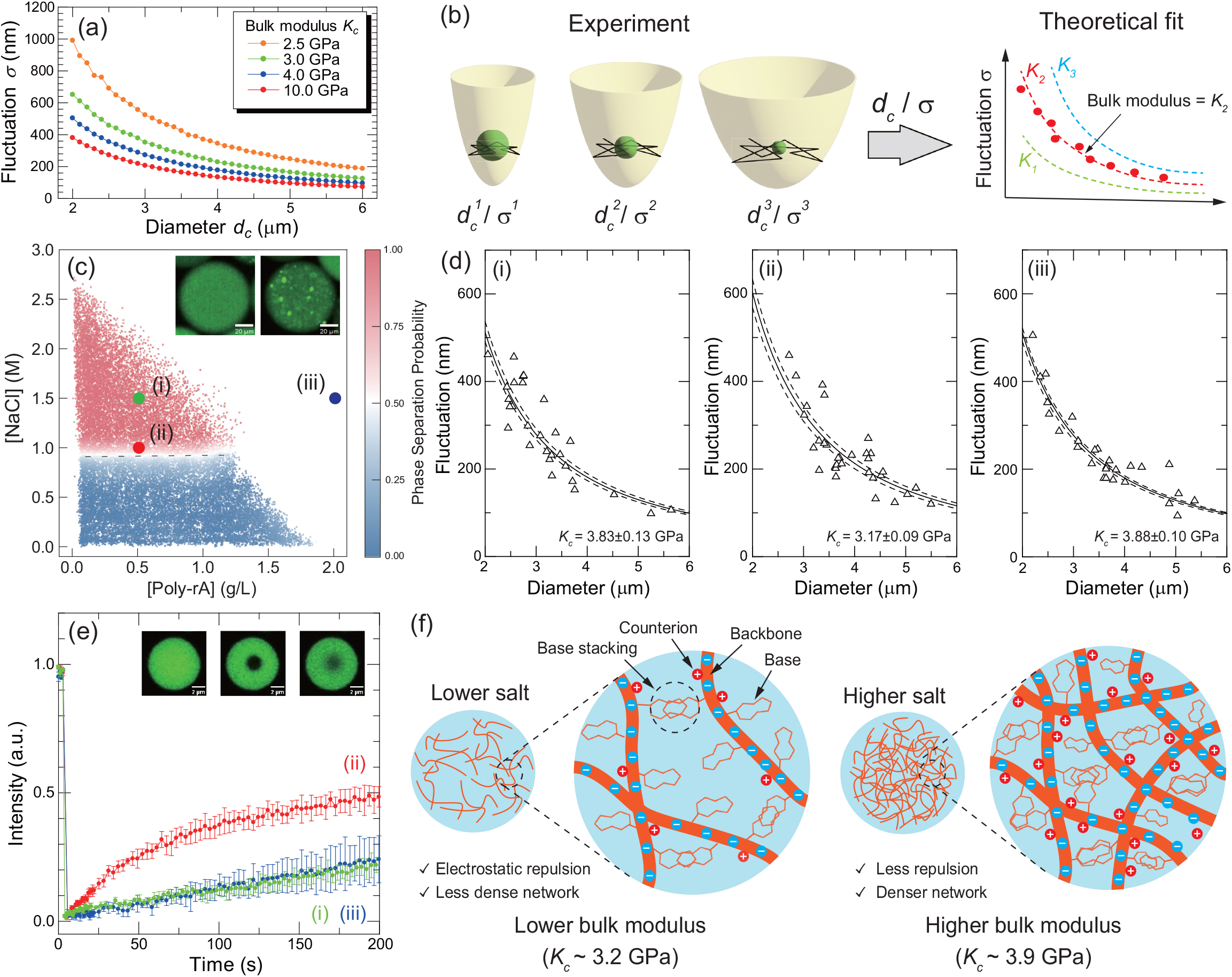
Change in the bulk modulus of poly-rA condensates under different environments. (a) The dependency of the fluctuation of condensates with various bulk moduli on the diameter of condensates under acoustic fields with *A*_*x,y*_ = 100 kPa and *f*_*x,y*_ = 35.2*/*35.3 MHz. (b) Workflow to analyze the bulk moduli of biopolymer condensates using acoustic trapping. (c) High-resolution phase diagram of poly-rA phase separation with various salt and polymer concentrations obtained using the droplet microfluidics platform (*n* = 30, 546). Insets show the fluorescent images of water-in-oil droplets without (left) and with phase separation (right). Scale bars: 20 µm. (d) Analysis on bulk moduli of poly-rA condensates formed under poly-rA and NaCl concentrations of (i) 0.5 g/L and 1.5 M (*n* = 28), (b) 0.5 g/L and 1.0 M (*n* = 32), and (c) 2.0 g/L and 1.5 M (*n* = 27), respectively. The solid and broken lines show the theoretical fitting lines of the average and 95% confidence interval of determined bulk moduli, respectively. The values in the panel denote mean±standard deviation. (e) FRAP curves of poly-rA condensates formed under various solution conditions. The insets show the fluorescence image of the condensate, formed with poly-rA and NaCl concentrations of 0.5 g/L and 1.0 M, immediately before (left) and after (middle) breaching of the dye and after 200 s (right), respectively. Scale bars: 2 µm. Error bars denote the standard deviation among three independent measurements. In panels (d) and (e), the numbers i-iii indicate the solution conditions shown in the phase diagram in panel c. (f) Schematic illustration of the interaction network within poly-rA condensates with various salt concentrations.

To determine the experimental conditions, we obtained a high-resolution phase diagram for the phase separation of poly-rA using a droplet microfluidic platform ^39,40^ (**Fig. 4c**). Briefly, we generated approximately 30,000 water-oil droplets with varying salt and polymer concentrations using a microfluidic device and examined which conditions led to poly-rA phase separation through microscopic observation (insets of **Fig. 4c**). Subsequently, poly-rA condensates formed under three different environments were trapped in acoustic potential wells, and their fluctuations in the acoustic potential well were observed. Finally, their bulk moduli were analyzed using Langevin dynamics simulations (**Fig. 4d**).

Under environmental conditions with poly-rA and NaCl concentrations of 0.5 g/L and 1.5 M (**Fig. 4d(i)**), respectively, the bulk modulus of the condensates was determined to be *K*_*c*_ = 3.83 ± 0.13 GPa (95% confident interval (CI) = [3.60, 4.11] GPa). When the NaCl concentration was reduced while keeping the poly-rA concentration constant (**Fig. 4d(ii)**), the bulk modulus decreased to *K*_*c*_ = 3.17 ± 0.09 GPa (95% CI = [3.03, 3.36] GPa). Increasing the poly-rA concentration to 2.0 g/L while maintaining the NaCl concentration at 1.5 M (**Fig. 4d(iii)**) resulted in a bulk modulus of *K*_*c*_ = 3.88 ±0.10 GPa (95% CI = [3.72, 4.10] GPa), which remained unchanged compared to the lower poly-rA concentration. These results indicate that the bulk modulus of poly-rA condensates is sensitive to changes in NaCl concentration but remains unaffected by changes in poly-rA concentration.

As shown in the phase diagram (**Fig. 4c**), the binodal phase boundary is parallel to the poly-rA concentration axis, indicating that the phase separation behavior of poly-rA primarily depends on the NaCl concentration. We also investigated the fluidity of poly-rA molecules within condensates using the fluorescence recovery after photobleaching (FRAP) ^41^ assay (**Fig. 4e**). Condensates formed under 1.5 M NaCl exhibited similar recovery curves regardless of the poly-rA concentration, with approximately 25% of the fluorescence recovering after 200 seconds. In contrast, condensates formed under lower NaCl concentrations demonstrated a more liquid-like nature, as indicated by a higher recovery rate in the FRAP curve. These results explain that poly-rA condensates exhibiting a more liquid-like nature possess lower bulk moduli.

## 3 Discussion

In soft matters, such as gels and rubbers, it is well-known that the bulk modulus is significantly larger than the tensile and shear moduli. The bulk modulus is typically 1 GPa, while the tensile and shear moduli generally range from 10 kPa to 100 kPa ^42^. This disparity arises from the different mechanisms underlying the restoring forces against compression and tensile deformation. Under tensile stress, the restoring force in rubbers primarily comes from entropic effects, where polymer molecules resist stretching to maintain their chain configurations to the greatest extent possible. In contrast, the restoring force against compression mainly results from short-range intermolecular interactions between interacting groups ^43^. Therefore, the denser the interaction network within soft materials, the higher the bulk modulus.

Our experimental results indicate that poly-rA condensates formed at higher salt concentrations exhibit higher bulk moduli than those formed at lower salt concentrations. Previous research has shown that the phase separation of poly-rA molecules is primarily driven by intermolecular base stacking ^44^. At higher salt concentrations, the intermolecular electrostatic repulsion, which originates from the negative charges on the backbone phosphate groups, is reduced due to electrostatic shielding by counterions (**Fig. 4f**). This reduction in repulsion facilitates the formation of a denser base-stacking network within the condensates, resulting in an increased bulk modulus (**Fig. 4f**). Therefore, measuring the bulk modulus of biomolecular condensates provides valuable insights into the interaction networks within these structures.

The storage elastic moduli, *G*′, of liquid-like biopolymer condensates have been reported to range from *G*′ ∼ 0.1 − 10 Pa ^14,15^, significantly lower than typical gels or rubbers. In such systems, macroscopic deformation is primarily governed by surface tension rather than elasticity, making accurate measurements of their elastic properties inherently challenging. Thus, utilizing acoustic trapping to measure the bulk modulus presents a novel approach for profiling the elastic behavior of mesoscale condensates at the single-particle level. This method provides a novel framework for enhancing our understanding of the interaction networks within condensates, offering deeper insights into the mechanism underlying condensate function and malfunction.

## 4 Conclusion

In this study, we demonstrated that acoustic trapping can be used to profile the mechanical properties of biopolymer condensates with precision and without physical contact. Our results showed that the 2D acoustic field generated by SAWs with frequencies of ∼ 50 MHz efficiently trapped and aligned poly-rA condensates based on the acoustic radiation force in a contactless manner. We presented the novel framework for non-invasive measurements of the mechanical properties of condensates by using the positional fluctuation of acoustically trapped condensates as an accurate probe of their bulk modulus. The framework could detect changes in the bulk moduli of poly-rA condensates as a consequence of changes in interaction networks within the condensates in response to environmental shifts. This novel approach not only provides a powerful tool for probing the mechanics of soft colloids at the single-particle level but also holds significant potential for applications across biology, materials science, and other disciplines.

## Materials and Methods

### Preparation of poly-rA condensates

Lyophilized poly-rA with a chain length of 2, 100-10, 000 nucleotides was purchased from Merck (1010862001). The lyophilized poly-rA was dissolved into nuclease-free water (AM9937, Thermo Fisher Scientific) to be the concentration of ∼10.0 g/L, determined by an absorbance measurement. Then, the poly-rA solution was mixed with HEPES buffer (pH 7.0) and poly-uracil labeled with cyanine-5 to be the final concentrations of 50 mM and 10 µg/mL, respectively. After mixing them well, the phase separation was induced by adding NaCl.

### Fabrication of piezoelectric chips for acoustic tweezers

Four IDTs were fabricated on a piezoelectric 128^°^ Y-cut LN substrate with a thickness of 0.35 mm. An Al thin film with a thickness of ∼300 nm was formed on the LN substrate by means of radio-frequency magnetron sputtering. A photoresist (AZ P4210, Merck) layer was spin-coated on the Al layer and exposed to UV light using a photomask. The pattern of the IDTs was formed by developing the photoresist layer (AZ developer, Merck) and subsequently removing the unnecessary Al thin film by chemical etching (Mixed acid Al etchant, Kanto Chemical Co., Inc.). After removing the remaining photoresist on the IDTs by a remover reagent (AZ Remover100, Merck), a silicon dioxide layer with a thickness of ∼200 nm was formed on the substrate for protecting the IDTs.

The fabricated LN chip possesses two pairs of IDTs, which are orthogonal to each other. The direction of each IDT is 45^°^ rotated from the X axis of the crystal because the 128^°^ Y-cut LN shows an anisotropy of a sound velocity of the Rayleigh wave in a plane. Consequently, we obtain the four equivalent IDTs with the sound velocity of the Rayleigh wave of *v*_*R*_ = 3, 590 m/s ^45^. The width of the electrode and the gap between the electrodes were designed using this sound velocity.

### Fabrication of microfluidic channels

The microfluidic channel was designed using AutoCAD and printed on the acetate film photomask (Micro Lithography). The SU-8 photoresist (SU-8 3050, KAYAKU Advanced Materials Inc.) was spin-coated on a polished silicon wafer. After baking the silicon wafer at 95 ^°^C for 20 min, the photomask was put on the top of the wafer, and then, the wafer was exposed to UV light. The microfluidic pattern was developed by soaking the wafer in propylene glycol methyl ether acetate (Sigma Aldrich). After drying and washing the SU-8 mold, a mixture of the PDMS elastomer and crosslinking agent (Sylgard184, Dow Corning) was poured on the SU-8 mold and incubated at 65 ^°^C for 3 h for polymerization of PDMS. The PDMS channel was cut from the mold using a scalpel and used for the experiments.

### Acoustic trapping experiments

The fabricated LN chip and PDMS microchannels were washed by isopropanol using a sonication bath for 5 min and dried at 65 ^°^C for 15 min. The LN chip and PDMS microchannels were assembled under a microscope for alignment. The surface of the chip and microchannel were treated by oxygen plasma (500 s; 80% power, Femto, Diener Electronics). The microchannel was filled with 1%(w/v) polyvinyl alcohol (PVA) solution for 15 min, and then, the PVA solution was removed by incubating the device on a hotplate with 115 ^°^C for 1 hour. After that, the channel was thoroughly washed by flushing ultrapure water for 15 min. These procedures are performed to make the surface of the microchannel hydrophilic ^46^ to prevent poly-rA condensates from the channel surface. The sample solution was introduced inside the microfluidic channel using the syringe pump (Cetoni, Nemesys S) with the flow rate of 100 µL/h. The alternative voltage whose frequency matches the resonant frequency of the IDTs was generated using a two-channel function generator (RS components, RSDG 6022X) and applied to the IDTs using needle-type contact probes (Harwin, P25-0423).

### Tracking analysis of condensate trajectory

Poly-rA condensates were visualized by a fluorescence microscope (CAIRN, openFrame Microscope) using a 40x objective lens (Nikon, Plan Fluor 40x). For the measurement of fluctuation of the Brownian motion, 1000 fluorescence images of identical condensates were sequentially acquired with the exposure time and frame rate of 10 ms and 10 fps, respectively. The acquired images were processed using the Trackmate algorithm ^47^, a plug-in of ImageJ software to acquire the trajectory of the Brownian motion. The diameters of the condensate were determined from the area of fluorescent images of the condensate.

To calculate the fluctuation from the trajectory, we determined one axis *a*, which minimizes the standard deviation of the distribution of the condensate position along that axis, and another axis *b*, which is perpendicular to *a* axis. We calculated the standard deviations along two axes, *σ*_*a*_ and *σ*_*b*_. Then, the averaged fluctuation *σ* was obtained as 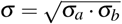. This operation corresponds to converting an ellipse with a minor and major axis of *σ*_*a*_ and *σ*_*b*_ into a circle with the equivalent area.

### Langevin dynamics simulation

For the Langevin dynamics simulation of the 2D Brownian motion of the condensate in an acoustic potential well, we used governing equations, **Eqs. 1,2,5,6** and **7**, and the following parameters: *T* = 298.15 K, *k*_*B*_ = 1.38 × 10^−23^ J/K, *ρ*_*m*_ = 997.1 kg/m^3^, *K*_*m*_ = 2.22 GPa, and *η*_*m*_ = 8.94 × 10^−4^ Pa·s. The simulation was performed by converting **Eq. 7** into a discrete form with a time step and spatial resolution of 100 µs and 10 nm using the Matlab software (version R2023b).

### Determination of bulk moduli of poly-rA condensates

The poly-rA condensates were trapped in the acoustic field with the frequencies and pressures of *f*_*x,y*_ = 35.2*/*35.3 MHz and *A*_*x,y*_ = 100 kPa, respectively. After the equilibrium of background flow, the fluctuations of the Brownian motion of ∼ 30 condensates with various diameters were recorded. The experimental data were compared to results from Langevin dynamics simulations conducted with a range of bulk moduli. Mean square errors (MSEs) were calculated between the experimental and simulated data. The bulk modulus that minimized the MSE was identified as the bulk modulus of the poly-rA condensates. To estimate the uncertainty of the determined bulk modulus, the bootstrap method was employed with the sampling number and iteration of 20 and 50, 000, respectively.

### High-resolution phase diagram obtained by PhaseScan

The microfluidic devices for acquiring the high-resolution phase diagram were fabricated by casting PDMS (Sylgard 184 kit; Dow Corning) on a master wafer, curing it at 65 ^°^C for 60 min, peeling it off, punching the holes for inlets and outlets, and bonding it to a 1-mm-thick glass slide (Epredia) after oxygen plasma activation in plasma oven (Diener Femto, 40% power for 30 s). Subsequently, hydrophobic treatment of the channels of the microfluidic device was performed. The channels were filled with 1% v/v trichloro(1H,1H,2H,2H-perfluorooctyl)silane (Sigma-Aldrich) in HFE-7500 fluorinated oil (3M™ Novec™ Engineered fluid) solution and incubated for 1–2 min. After the incubation, channels were washed with HFE-7500 fluorinated oil and dried under airflow.

Phase diagrams were constructed using the semi-automated microfluidic platform ‘PhaseScan’ ^39,40^. Briefly, poly-rA and NaCl stock solutions were mixed at various mass ratios and encapsulated in water-in-oil droplets and individual microenvironments using a microfluidic device. Fluorescence images of droplets in the observation chamber of the microfluidic device were taken using an epifluorescence microscope (Cairn Research) equipped with 10× objective (Nikon CFI Plan Fluor 10×, NA 0.3) and analyzed via automated image analysis script to detect condensates and to determine peptide and RNA concentrations in individual droplets. The data was plotted as a color-coded scatter plot, where each point represents an individual microfluidic droplet, and color represents the phase separation probability (1 or dark red – phase separated, 0 or dark blue–mixed) assigned to the droplet. This probability was calculated by averaging the phase state assignment of neighboring droplets. The radius of the neighborhood was defined as a percentage of data range (5%).

### Fluorescence recovery after photobleaching (FRAP) assay

Fluorescence recovery after photobleaching. FRAP (fluorescent recovery after photobleaching) was performed using the Stellaris confocal microscope equipped with a 63×oil immersion objective (Leica HC PL APO 63×/1.40 Oil CS2, NA 1.4). A 488 nm argon laser at 100% power was used to bleach a disk-shaped 1 µm^2^ area in the center of condensates. The half-lives of fluorescence recovery, tau, were obtained by analyzing FRAP kymographs using the built-in software. We report the average and standard deviation of 3 repeats for each condition.

## Author Contributions

K.N., T.S., and T.P.J.K. designed and managed this research project. K.N., T.S., and L.L.G. established the experimental setup and protocols. K.N., T.S., and N.A.E. acquired experimental data and analyzed them. K.N., T.S., L.L.G., N.A.E., H.O., and T.P.J.K. discussed the experimental results and wrote the manuscript.

## Acknowledgements

This study was supported by the Japan Society for the Promotion of Science (JP22K14013; K.N.), JKA and its promotion funds from AUTORACE (2024M-583; K.N.), the National Institutes of Health Oxford-Cambridge Scholars Program (L.L.G.), the Cambridge Trust’s Cambridge International Scholarship (L.L.G.), the Intramural Research Program of the NIH, The National Institute of Diabetes and Digestive and Kidney Diseases (NIDDK)(L.L.G), the Royall scholarship (N.A.E), the European Union’s Horizon 2020 research and innovation programme under the Marie Skłodowska-Curie grant MicroREvolution (agreement no. 101023060; T.S.), the Frances and Augustus Newman Foundation (T.S.), and the ERC grant DiProPhys (agreement ID 10100161; T.P.J.K.).

