## Supplementary Information for "Mechanical Profiling of Biopolymer Condensates through Acoustic Trapping"

### Table of Content

**Supplementary Result: Page S2-4**
**Captions of Supplementary Movies: Page S5**

**Supplementary Figures**

**1. Appearance of the acoustic device**

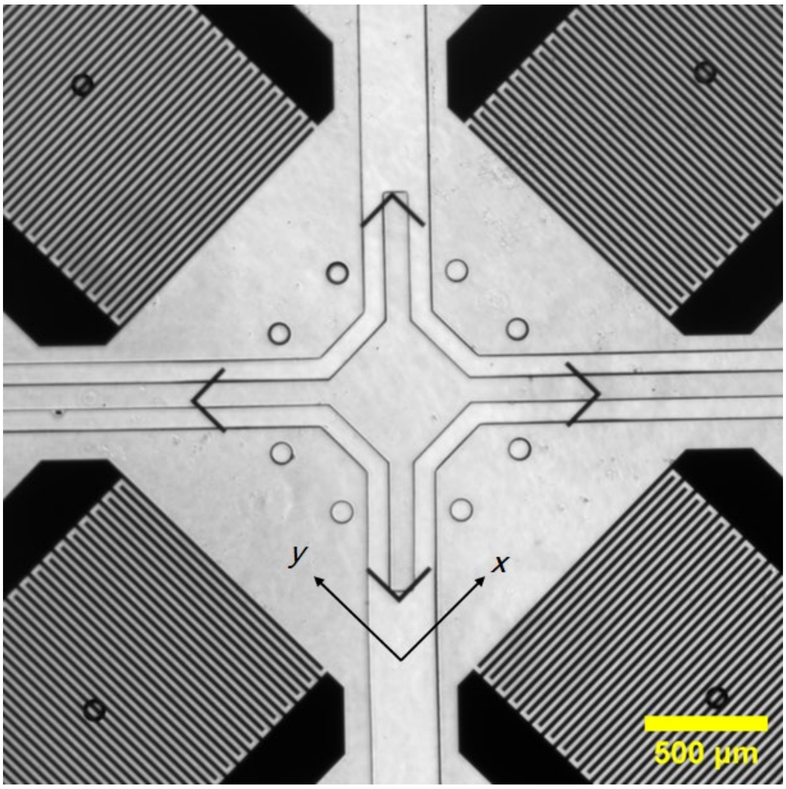


**Figure S1.** The micrograph of the acoustic device. Scale bar: 500 μm.

### 2. Lattice structure of polystyrene particle clusters under the 2D acoustic field

**Figure S2a** illustrates the 2D acoustic potential created by standing pressure fields with frequencies of $f_{x,y}$ = 70.4/70.5 MHz. In the calculations, we assumed phase velocities of $v_R^x = v_R^y$ =3,590 m/s[(1)] for the Rayleigh waves propagating on a 128° Y-cut LN substrate, oriented at a 45° angle from the X-axis of the crystal, as the IDTs were fabricated in this direction in the experiments. The resulting acoustic potential exhibits a 2D lattice structure with lattice constants along the x- and y-axes of $a_{x,y} = \frac{v_R^{x,y}}{2f_{x,y}} \sim$ 25.5 μm. We experimentally verified the formation of this acoustic potential using polystyrene (PS) particles with an average diameter of 1.1 μm (**Fig. S2b** and **Movie S1**). The randomly dispersed PS particles migrated to the nearest trapping points immediately after generating SAWs with frequencies of $f_{x,y}$ = 70.4/70.5 MHz. After 10 seconds, the particle clusters formed the 2D lattice structure with lattice constants of $a_{x,y}$ = 25.3 ± 1.0/25.6 ± 1.9 μm, which is consistent with the theoretical values. Then, we apply this experimental setup in the following experiments to study the mechanical properties of biopolymer condensates.

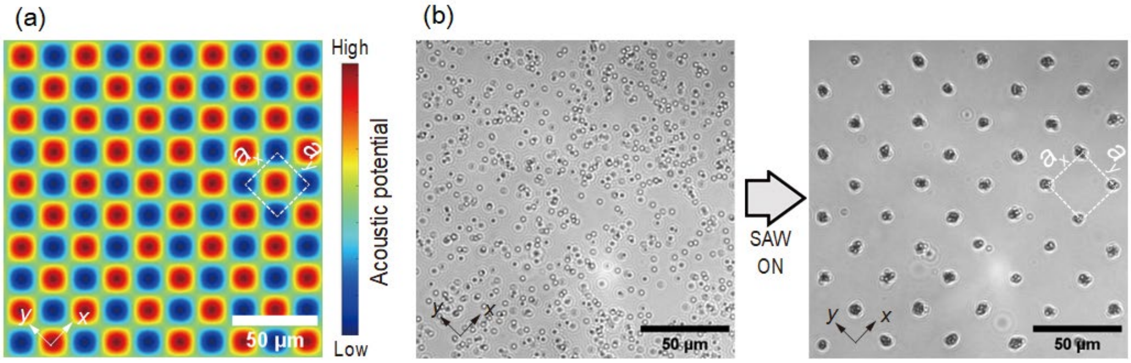


**Figure S2.** (a) Theoretical calculation of the acoustic potential formed by the 2D standing pressure field with $f_{x,y}$ = 70.4/70.5 MHz. (b) Formation of the 2D lattice structure of 1.0-μm polystyrene particle clusters by SAWs with $f_{x,y}$ = 70.4/70.5 MHz. Scale bars: 50 μm.

**3. Calibration of the acoustic pressures:** To estimate the relationship between the applied voltage on the interdigital transducers and acoustic pressures created in the microfluidic chamber, we performed the calibration experiment using silica particles with diameters of 2.05±0.07 and 3.04±0.14 μm. We measured the fluctuation of the Brownian motion of silica particles suspended in water under seven various voltages, as shown by the plots in Fig. S3. We also conducted the Langevin dynamics simulation using parameters for silica particles ($\rho_c = 2.65 \times 10^3$ kg/m³ and $K_c = 36.5$ GPa) and water ($\rho_m = 0.997 \times 10^3$ kg/m³ and $K_m = 2.2$ GPa). By fitting the experimental data with the simulation results by varying the acoustic pressures as a parameter, we obtained $A_{x,y} = 100$ kPa for the applied voltage in the experiment of 5 Vpp. This acoustic pressure values were used for analyzing the bulk moduli of poly-rA condensates in **Fig. 4d** in the main text.

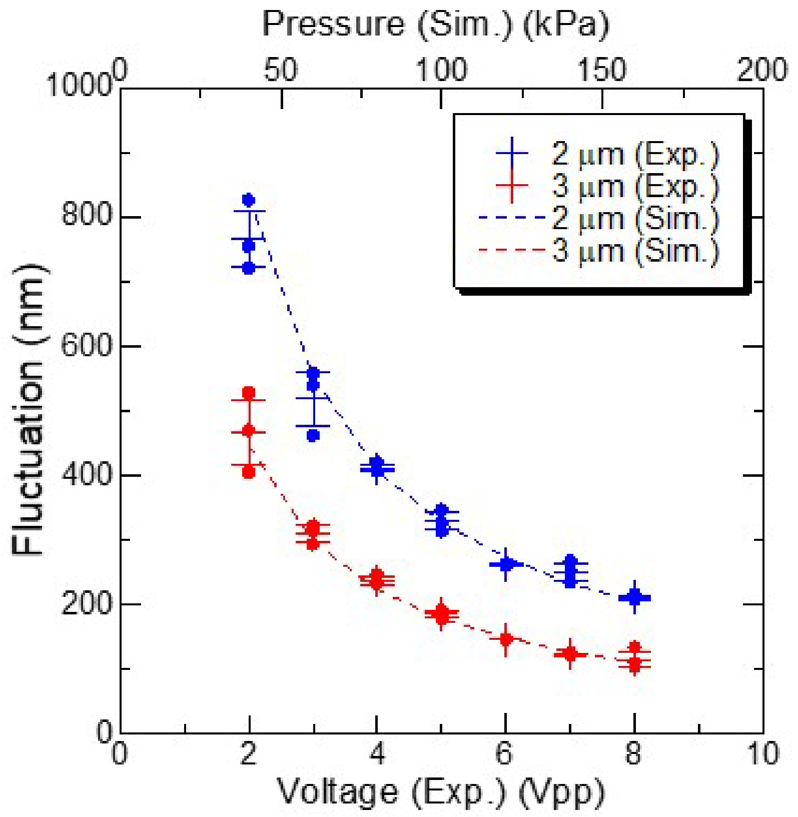


**Figure S3.** Fluctuation of silica particles with various applied voltages on the interdigital electrodes in the experiments and fitted simulation results. Error bars denote standard deviation among three independent measurements.

**Captions of Supplementary Movies**

**Movie S1:** Formation of the two-dimensional lattice structure of polystyrene particle clusters under the acoustic field with frequencies of $f_{x,y}$ = 70.4/70.5 MHz. The average diameter of polystyrene particles is 1.0 μm. The acoustic field was turned on at $t$ = 12.0 s. Scale bar: 50 μm.

**Movie S2:** Acoustic trapping of poly-rA condensates under acoustic field with frequencies of 49.6/49.7 MHz. The acoustic field was turned on at $t$ = 2.5 s. The condensates are formed with [poly-rA] = 1.0 mg/mL, [NaCl] = 1.0 M, and [HEPES] = 50 mM. Scale bar: 10 μm.

**Movie S3:** Acoustically induced coalescence of poly-rA condensates. The acoustic field was turned on at $t$ = 0.2 s. The condensates are formed with [poly-rA] = 1.0 mg/mL, [NaCl] = 1.0 M, and [HEPES] = 50 mM. Scale bar: 5 μm.
